# A new cloud forest tree *Lychnodiscus bali* (Sapindaceae), Critically Endangered from the Bali-Ngemba Forest reserve, N.W. Region, Cameroon

**DOI:** 10.1101/2023.11.04.565640

**Authors:** Martin Cheek, Jean Michel Onana, Olivier Lachenaud

**Affiliations:** Royal Botanic Gardens, Kew, Richmond, Surrey, U.K.; University of Yaoundé I, Faculty of Science, Department of Plant Biology P.O Box 812 Yaoundé, Cameroon; IRAD-National Herbarium of Cameroon, PO Box 1601 Yaoundé, Cameroon; Evolutionary Biology and Ecology, CP 160/12, Faculté des Sciences, Université Libre de Bruxelles, 50 Avenue F. Roosevelt, BE-1050 Bruxelles, Belgium; National Botanic Garden of Belgium, Domein van Bouchout, BE-1860 Meise, Belgium

**Keywords:** Bamenda Highlands, Biodiversity crisis, Extinction, *Lychnodiscus*, Tropical Important Plant Areas

## Abstract

We describe *Lychnodiscus bali* (Sapindaceae) a new species to science, from the Bali Ngemba Forest Reserve of NW Region Cameroon, the last major remnant of cloud forest in the Bamenda Highlands of Cameroon, recently evidenced as a Tropical Important Plant Area (TIPA or IPA). Confined on current evidence to upper submontane forest, the species is threatened by expanding habitat clearance for farms and is assessed as Critically Endangered. A small tree, attaining 3 *–* 4 m height, it is the first new species to be added to this Guineo-Congolian tree genus in 50 years, the third recorded from Cameroon, and takes the number of species in the genus to eight. It has the highest known altitudinal range (1700 – 1950 m alt.), of any species of the genus.

Previously identified as *Lychnodiscus grandifolius*, the new species differs in the shorter length of the distal leaflets (12 – 18 cm vs 22 – 39 cm long), in the abaxial surface lacking glands (vs glands flat and conspicuous), tertiary nerves hairy (vs glabrous), flowers at anthesis 8 – 11 mm long (vs 5 – 7 mm long).

*Lychnodiscus bali* is described, illustrated and its extinction risk assessment as Critically Endangered is presented. We discuss its discovery in the context of other recently discovered and highly threatened or even extinct plant species in the Cameroon Highlands, and the importance of their conservation.

We present an updated key to the identification of the species of the genus,and discuss its classification in the context of recent molecular phylogenomic studies. Previously placed in Cupanieae by Radlkofer, the authors contend that *Lychnodiscus* should now be placed in the reconstituted Nepheliaeae in the revised 2021 intrafamilial classification of Buerki et al., probably close to the genera *Aporrhiza* and *Laccodiscus*. However, until the genus is included in molecular studies this cannot be confirmed and its sister relationship remains speculative.

## Introduction

The genus *Lychnodiscus* Radlk. (Sapindaceae) occurs in tropical Africa and is largely restricted to the Guineo-Congolian region, from the Republic of Guinea (Gosline et al. 2023a, 2023b) to S. Sudan (Darbyshire et al. 2015) and southwards to Uganda (Davies & Verdcourt 1998) and DR Congo (Hauman 1960). It is easily recognised, among African Sapindaceae, by its actinomorphic flowers (many other African genera of the family being zygomorphic) in which there are two concentric discs, often resembling a cup and saucer (in other genera there is usually no more than one disc). The five sepals are quincuncially imbricate, united at the base. The ovary is 3-locular with a distinct style, the locules uniovulate. The fruit is dehiscent, the inside of the three leathery valves being red or pink and glabrous, and the three large seeds entirely covered in a glossy, sometimes sticky, red or orange aril or seedcoat so that we conjecture that they are likely dispersed by primates.

*Lychnodiscus* has been placed until recently in the standard tribal classification of the family (Radlkofer 1932) in the pantropical tribe Cupanieae Rchb., which has 40 genera, of which only five are in Africa (Fouilloy & Hallé 1973): *Aporrhiza* Radlk., *Laccodiscus* Radlk., *Lychnodiscus, Eriocoelum* Hook.f., and *Blighia* K.D.Koenig. The tribe was characterised by uniovulate locules, paripinnate leaves (although saplings of *Lychnodiscus* and *Laccodiscus* often have simple leaves) and loculicidally dehiscent fruit, often with fleshy valves (Radlkofer 1932). The flowers are usually regular, the petals with an adaxial scale that is bifid or united to the limb to form a funnel (Fouilloy & Hallé 1973). *Lychnodiscus* has not yet been included in any published molecular phylogenomic studies (Buerki et al. 2021; Joyce et al. 2023). However, these studies, based on over 300 nuclear genes, indicate that the similarities between these five African genera of Radlkofer’s Cupanieae are due to convergence since they are scattered among genera of other of Radlkofer’s tribes in phylogenetic trees which have high support. Thus, Cupanieae in the sense of Radlkofer (1932) is polyphyletic. Realignment of tribes in Sapindaceae was provided by Buerki et al. (2021). We conjecture that the closest relatives of *Lychnodiscus* include probably *Aporrhiza* and *Laccodiscus*, both of which differ by their simple disk and bicoloured seeds (orange at base, black at apex). The former, which shares with *Lychnodiscus* the very unusual character of a radicle opposite the hilum, may also be separated by its 2-locular ovary and valvate sepals, while the latter has fruit valves hirsute inside, an inferior radicle, petals lacking an internal scale (but with lateral appendages) and leaves often with pseudostipular lower leaflets. *Aporrhiza* and *Laccodiscus* are in the same subclade (with non Cupanieae genera *Pancovia* Willd., *Placodiscus* Radlk., and *Haplocoelopsis* F.G.Davies) of Buerki et al. (2021) and Joyce et al. (2023). This subclade, all genera being tropical African, is part of clade 9, comprising the redelimited tribe Nephelieae of Buerki et al. (2021), currently with 16 genera including 116 species. However, as newly reconstituted, there is as yet no morphological delimitation of this tribe available: “We have not yet identified any morphological synapomorphies that define this clade” (Buerki et al. 2021). We consider, *Lychnodiscus* is likely to be placed in Nephelieae with these genera when eventually the genus is included in molecular studies.

Like many genera of African Sapindaceae, *Lychnodiscus* has not been revised as a whole since Radlkofer (1932), although regional treatments have been published for W. Africa (Keay 1958), Cameroon and Gabon (Fouilloy & Hallé 1973a, 1973b), DR Congo, Rwanda and Burundi (Hauman 1960) and East Africa (Davies & Verdcourt 1998). Seven species of *Lychnodiscus* are currently accepted (Lebrun & Stork 1992; African Plant Database, continuously updated, POWO continuously updated): *L. brevibracteatus* Fouilloy, *L. cerospermus* Radlk. (with three varieties), *L. dananensis* Aubrév. & Pellegr., *L. grandifolius* Radlk., *L. multinervis* Radlk., *L. papillosus* Radlk., and *L. reticulatus* Radlk. However, the number of species is probably under-estimated and the genus is much in need of a new revision. Two species are recorded from W. Africa (Keay 1958), two from Cameroon (Fouilloy & Hallé 1973a), two from Gabon (Fouilloy & Hallé 1973b, Lachenaud et al. 2018) and two also from D.R.Congo (Hauman 1960). The genus is most taxon-diverse in DR Congo, with four taxa (Hauman 1960). However, the genus is under-researched, for example until recently (Lachenaud et al. 2018), of the eight specimens of *Lychnodiscus* listed for Gabon (Sosef et al. 2006) only one was identified to species.

The species are shrubs to medium-sized trees and occur mostly in lowland forest, *Lychnodiscus cerospermus* is unusual in that it is found in the understorey of forest at 1000– 1500 m in altitude which is often inundated part of the year (Bosch 2012).

*Lychnodiscus* species are usually infrequent within their ranges and occur as isolated individuals (although *L. grandifolius* may be locally frequent in Gabon, Lachenaud pers. obs.). Eilu *et al*. (2004) in their study of four forests in the Albertine Rift of Uganda, enumerated 212 tree species of diameter above 10 cm at 1.5 m above ground, assessing the rarity status of each species; *Lychnodiscus cerospermus* was among the four rarest of those 212 species, having both a restricted range and low population density.

Few species have recorded uses apart from firewood (Burkill 2000). However, the wood of *Lychnodiscus cerospermus* is used in DR Congo for construction and to make mortars (Bosch 2012). Young leaves of the same species are consumed by chimpanzees in the Budongo Forest of western Uganda but are not a major part of their diet (Bates 2005), perhaps because of the rarity of these trees.

In connection with the Cameroon Tropical Important Plant Areas programme (Darbyshire *et al*. 2017; Murphy et al. 2022; Murphy et al. 2023), in May 2022, two specimens from the Bali Ngemba Forest Reserve, *Ghogue* 1080 and *Cheek* 10503 (both K!) that had been identified in Harvey et al. (2004) as *Lychnodiscus grandifolius* Radlk. were reviewed. Although previously unambiguously identified as this species, the specimens had been annotated “Possibly a montane form, 6 not 5-jugate, leaves narrower, hairy on midrib and veinlets, rhachis - not glabrous” by the first author. The third author, in the course of reviewing *Lychnodiscus grandifolius* in the context of species range extensions into Gabon, had later added the annotation “Presumably sp. nov. differs from *grandifolius* by leaves sparsely pubescent beneath and lacking glandular dots; flowers larger; calyx and ovary with long golden-brown indumentum (vs. very short indumentum); montane habitat”. In Lachenaud et al. (2018) it is stated: “Collections from the western Cameroon highlands, previously identified as *L. grandifolius* (e.g. Cheek et al. 2004: 399) appear to represent two new taxa. The first of these, from Bali Ngemba F.R. (*Cheek* 10503; *Ghogue* 1080) differs from *L. grandifolius* by the longer indumentum of the leaflets, inflorescences and ovaries, the absence of leaf glands, and the larger flowers. The second taxon, from Mt Manenguba, is only represented by a poor fruiting specimen (*D*.*W. Thomas* 3110) which differs from *L. grandifolius* by the larger size and dark brown indumentum of the fruits.” Further research showed additional characters separating the Bali Ngemba high altitude taxon from the lowland *L. grandifolius* (Table 1) and the former is here described as a new species, *Lychnodiscus bali*. The Manenguba taxon referred to is not addressed further here, being only represented by incomplete fruiting material, and further collections not being possible due to the ongoing warfare in NW and SW Regions of Cameroon.

**Table 1.**
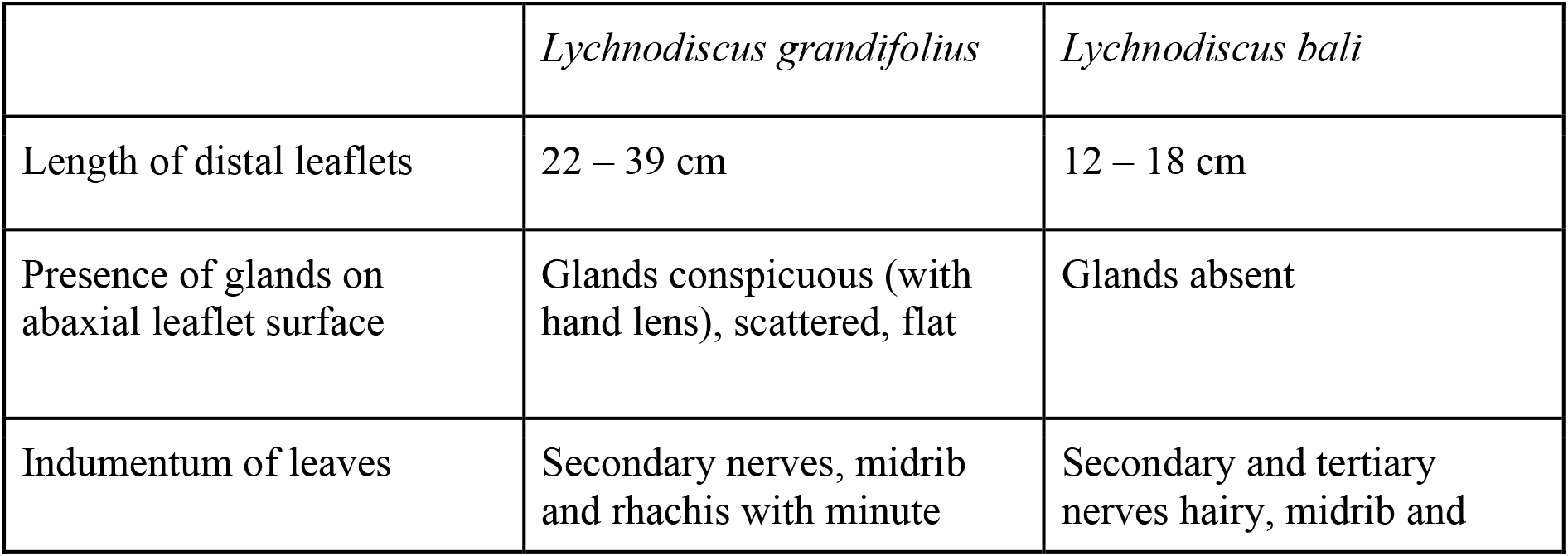
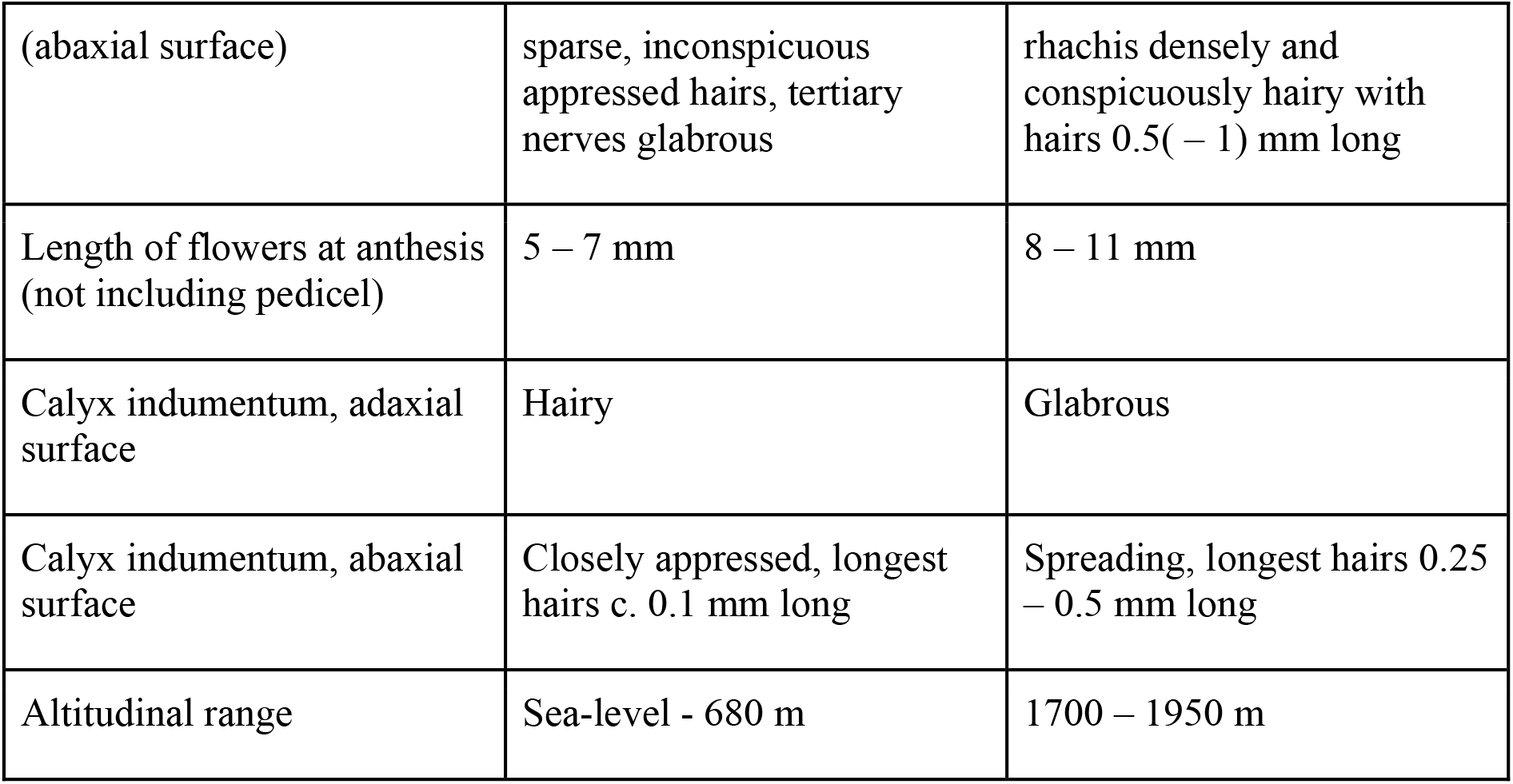
Characters separating *Lychnodiscus grandifolius* from *L. bali*.

## Materials and methods

Specimens were collected using the patrol method and processed as documented in Cable & Cheek (1997). Identifications were made following Cheek (2023). Herbarium citations follow Index Herbariorum (Thiers et al., 2020). All specimens were seen by one or more authors. Specimens of *Lychnodiscus* were studied at BR, BRLU, K, LBV, P, WAG, and YA. The National Herbarium of Cameroon, YA, was specially searched for additional material of the new taxon as was Tropicos (http://legacy.tropicos.org/SpecimenSearch.aspx). Images for WAG specimens, now at Naturalis, were studied at https://bioportal.naturalis.nl/?language=en and those from P at https://science.mnhn.fr/institution/mnhn/collection/p/item/search/form?lang=en_US. We also searched *JStor Global Plants* (*2020*) for additional type material of the genus not already represented at K.

Binomial authorities follow the International Plant Names Index (IPNI, 2020). The conservation assessment was made using the categories and criteria of IUCN (2012). Herbarium material was examined with a Leica Wild M8 dissecting binocular microscope fitted with an eyepiece graticule measuring in units of 0.025 mm at maximum magnification. The drawing was made with the same equipment using Leica 308700 camera lucida attachment. Flowers from herbarium specimens of the new species described below were soaked in warm water to rehydrate the flowers, allowing dissection, characterisation and measurement. The description follows the terms of Beentje & Cheek (2003), format and conventions of Cheek et al. (2021).

### Taxonomic Results

### Key to the species of *Lychnodiscus* (after Fouilloy & Hallé 1973)

1.Abaxial leaflet surface glabrous or occasionally with a few short hairs… 4

1. Abaxial leaflet surface pubescent, with hairs 0.5 – 0.5 mm long on the secondary nerves. 2

2. Leaflet margins dentate distally; petiole 5 – 6 cm long… ***L. papillosus***

2. Leaflet margins entire, or sometimes toothed at apex; petiole 10 – 17 cm long… 3

3. Lateral nerves 20 – 22 on each side of the midrib; hairs crisped; inflorescence c.30 cm long; petals 4 – 5 mm long… ***L. dananensis***

3. Lateral nerves 9 – 12 on each side of the midrib; hairs straight; inflorescence >60 cm long; petals 8 – 9 mm long… ***L. bali sp*.*nov***.

4. Leaflets entire; length: breadth c. 2:1; >20 – 25 cm long… 5

4. Leaflets denticulate (sometimes entire in *L. reticulatus*); length: breadth c. 3:1; 20 – 25 cm long… 6

5. Leaflets with 10 – 14 nerves on each side of the midrib, glabrous; stamens 7 – 9 …………………………………………………………………………….…***L. grandifolius***

5. Leaflets with 18 – 24 nerves on each side of the midrib, nerves red glandular hairy; stamens 10 – 12… ***L. multinervis***

6. Leaflets entire, or, near apex, with 3 – 4 teeth to c. 1 mm long; glabrous… 7

6. Leaflets dentate for the distal 2/3, 12 – 18 on each side, teeth c. 2 mm long; nerves sparsely hairy… ***L. cerospermus***

7. Leaflets 2 – 3 pairs; margin entire; with 5 – 6 nerves on each side of the midrib; petiolule c. 4 mm long… ***L. brevibracteolatus***

7. Leaflets 4 – 6 pairs; apex with 3 – 4 teeth; c. 5 – 6 nerves on each side of the midrib; petiolule 2 – 3 mm long ***L. reticulatus***

***Lychnodiscus bali*** Cheek *sp. nov*. Type: Cameroon, North West Region, Mezam Division, Bali Ngemba Forest Reserve, 5^0^ 48.87’ N, 10^0^ 0.5.56’ E, path from first valley to highest point of reserve, fl.fr. 13 Nov. 2000, *Cheek* 10503, with Biye, Nana, Iwanaka, Wanduku, Ach Nkankanu, van de Rheede, Garcia, Sam & Tadjouteu (holotype K000746446; isotypes MO, WAG0424201, YA). (Fig. 1).

**Fig 1.**
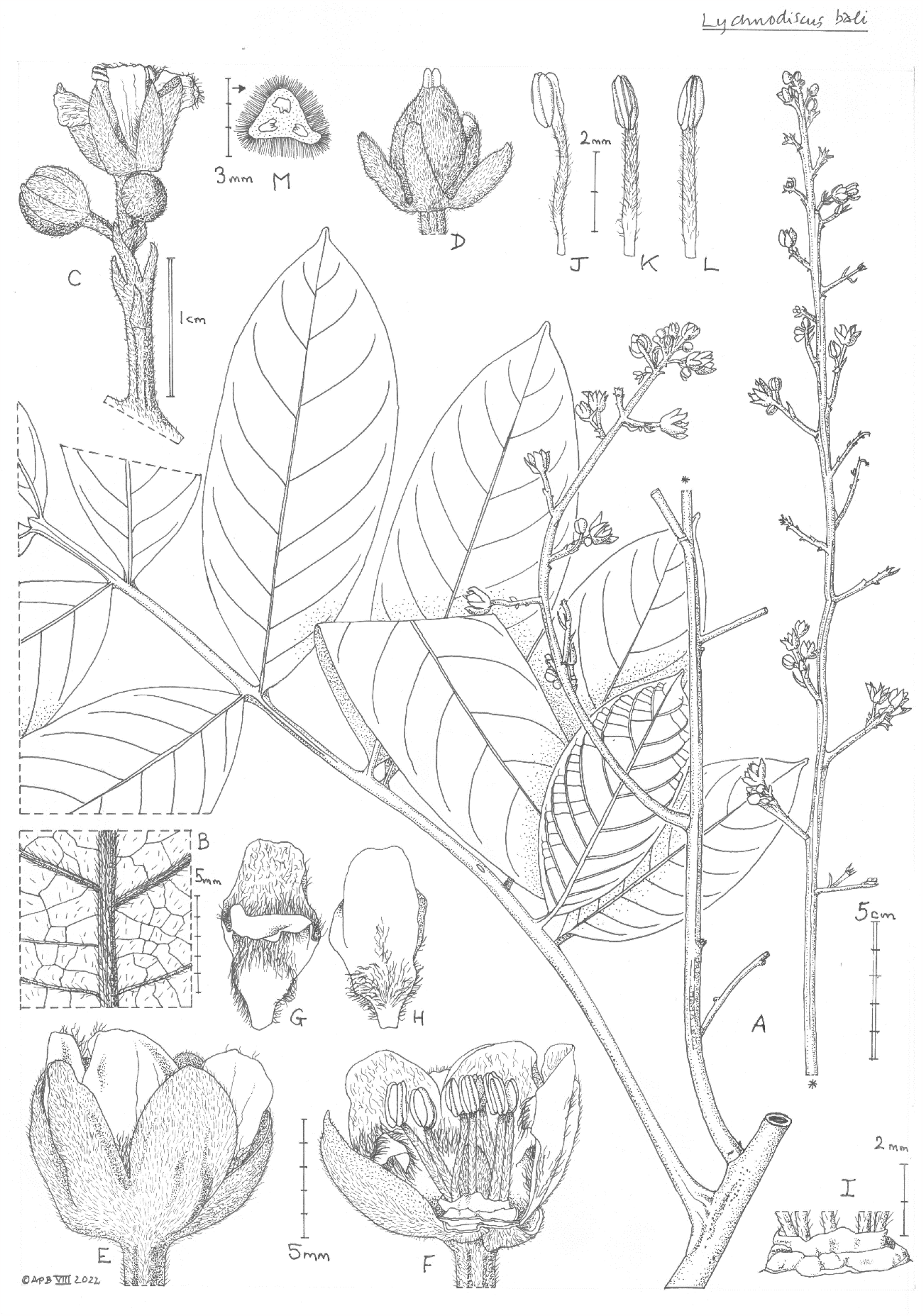
*Lychnodiscus bali* **A** habit, leaf, inflorescence and portion of stem from near stem apex; **B** abaxial leaf surface (detail); **C** partial-inflorescence showing partial-peduncle, bracts and flowers; **D** immature fruit; **E** flower, side view; **F** flower with proximal sepals and petals removed; **G** petal adaxial view; **H** petal abaxial view; **I** double disc and base of filaments; **J-L** stamens, respectively side, abaxial and adaxial views; **M** ovary transverse section. All drawn from *Ghogue* 1080 by ANDREW BROWN.

*Lychnodiscus grandifolius sensu* Cheek (2004:125) non Radlk.

*Shrub or tree* – 5 m tall, sparsely branched. *Stem* subterete 8 – 14 mm diam. at distal flowering nodes, minutely densely puberulent with inconspicuous simple brown patent hairs 0.025 – 0.75 mm long; lenticels dark brown, orbicular 0.25 – 0.75 mm. *Leaves* alternate, pinnately compound, 48 – 50 x 38 – 40 cm, 6-jugate, rhachis internodes plano-convex 3.5 – 6 cm long, indumentum as stem. *Leaflets* chartaceous, entire, discolorous, drying grey-black on adaxial surface and pale to mid brown on abaxial surface, opposite or subopposite, terminal leaflet vestigial 1 – 2 x 1 – 2 mm, median leaflets larger than proximal or distal. *Proximal leaflet* pair smallest, elliptic, 9.5 – 11.5 x 4 – 5.5 cm, apex obtuse or with a small acumen c. 2 x 2 mm, base acute, adaxial surface glabrous, midrib and lateral nerves impressed; abaxial surface with glands and domatia absent, lateral nerves c. 9 on each side of the midrib, tertiary nerves subscalariform, quaternary nerves reticulate, conspicuous, hairs patent, dull brown, simple 0.25 – 0.5 mm long, moderately dense on areolae, 0.5 (– 1) mm long on nerves, petiolules and leaf axis. *Median leaflet* pair as the proximal leaflets, but oblong-elliptic, 16.5 – 20 x 6.5 – 7.5 cm, apex obtuse to rounded, base obtuse to acute, slightly asymmetric, lateral nerves 10 – 12 pairs. *Terminal leaflet* pair obovate, 11.5 (– 18) x 5.8 cm, apex rounded, base acute, lateral nerves c. 10 pairs. *Petiolules* stout, subconical, dorsiventrally compressed, 4 – 5 mm long, 5 – 6 mm wide at base, 3 – 4 mm wide at apex, blade decurrent to the distal margins, densely hairy as stem. *Petiole* drying dark brown, terete (base thickened and decurrent), 11 – 11.5 x 0.4 – 0.6 cm, indumentum as stem. *Buds* supra-axillary, inserted c. 7 mm above the axil, globose, c.2 mm diam., densely short pubescent. *Inflorescence* terminal and axillary in distal nodes, thyrsoid, 60 – 66.5 x 22 – 28 cm, peduncle 9 – 12 x 0.6 cm; rhachis 48 – 57.5 x 0.4 – 0.5 cm, internodes 3.5 – 6.5 cm long, partial-inflorescences (secondary inflorescences) 24 – 31, most proximal partial-inflorescence 23 – 26.5 cm long each with 9 – 13 secondary partial-inflorescences each 1.5 – 3 cm long, 3 – 6-flowered; distal 20 – 25 partial-inflorescences 1.5 – 4 cm long; bracts linear-oblong, patent (4 –)5 – 6 x 1 mm; bracteoles as the bracts, 2 – 3 mm long, indumentum as stem, dense. *Pedicels* (3 –)5 – 6 x 1 – 1.2 mm, articulated at junction with partial-inflorescence, indumentum as stem.

*Flowers* hermaphrodite, actinomorphic, *Sepals* 5, imbricate in bud, splayed open at anthesis, dimorphic, oblong-ovate 6.5 – 7 x 4.5 mm, apex obtuse, outer and inner surface densely hairy, as pedicel, excepting the glabrous marginal 1 mm of both sides of the two inner sepals, and one side of the intermediate sepal. *Petals* 5, white in life, drying red-brown, slightly exserted from the sepals at anthesis, unguiculate, obovate or elliptic, 8 – 9 x 3 – 4 mm, apex broadly rounded, distal third to half of petal sparsely to moderately hairy on adaxial surface, glabrous adaxially or with a dew hairs along the midrib; inner appendage inserted c. 5.5 mm from the base, transversely oblong, as wide as petal, 1.5 mm long revolute, densely puberulent at the base in the axil with the main petal and on the oueter surface at the base, hairs translucent; the distal part of the appendage glabrous on both surfaces; proximal third to half of petal narrowing gradually towards base, adaxial surface glabrous, abaxial densely hairy; basal claw c. 2 mm long, 1 mm wide, glabrous on both surfaces. *Stamens* 8 – 9, free, equal, 6 – 7 mm long, filaments cylindrical, 4.5 – 5 x 0.4 mm, slightly curved or straight, with patent, translucent simple hairs 0.2 mm long, anther dull white, basifixed, more or less oblong, 1.8 x 1 x 0.75 mm, latrorse and introrse, dorsal surface with the two thecae separated by a broad connective, connective sometimes with a few hairs as the filament. *Discs* 2, closely surrounding the base of the ovary, outer disc 5 mm wide, inner disc c.4 mm wide, raised above the outer, both discs red, erect, thin, crenate c. 0.3 – 0.5 mm long. *Ovary* superior, 3-locular, trigonous-rounded, 5 x 4 mm, densely dull yellow hairy, hairs erect, 0.75 mm long. Style 3-lobed, stout, 2 x 1 mm, indumentum as ovary; stigma barely 3-lobed, glabrous. *Fruit and seed* not seen.

### RECOGNITION

Differs from *Lychnodiscus grandifolius* in the shorter length of the distal leaflets (12 – 18 cm vs 22 – 39 cm long), in the abaxial surface lacking glands (vs glands flat, sometimes red, and conspicuous), tertiary nerves hairy (vs glabrous), flowers at anthesis 8 – 11 mm long (vs 5 – 7 mm long). Additional diagnostic characters are given in Table 1.

### DISTRIBUTION & HABITAT

CAMEROON, N.W. Region, Bamenda Highlands, Bali Ngemba Forest Reserve 1700 – 1950 m alt., with *Hannoa ferruginea, Rhaphiostylis beninensis, Pentas ledermannii, Synsepalum brevipes, Rothmannia urcelliformis, Cyperus renschii* var. *renschii, Cuviera longiflora, Pseudognaphalium luteo-album, Adenia* cf. *cissampeloides, Ficus ardisioides* subsp. *camptoneura, Psychotria subpunctata, Amorphophallus staudtii, Stephania abyssinica* var. *abyssinica, Aframomum* sp. 1 of Bali Ngemba, *Pseudospondias microcarpa*. In disturbed areas (see Notes below).

### SPECIMENS

**CAMEROON**, N.W. Region, Bali Ngemba Forest Reserve, Mantum to Pigwin (sic.), 1950 m alt., fl. 11 Nov. 2000, *Ghogue* 1080 with Julia Garcia, Christo Van Der Rheede, Benedict Pollard, Emmanuel Sama, Petula Nabukeera, Francis Njie, (K barcode K000746448, P, YA); 5^0^ 48.87’ N, 10^0^ 0.5.56’ E, path from first valley to highest point of reserve, fl.fr. 13 Nov. 2000, *Cheek* 10503, with Biye, Nana, Iwanaka, Wanduku, Ach Nkankanu, van de Rheede, Garcia, Sam & Tadjouteu (holotype K000746446; isotypes MO, WAG0424201,YA).

### CONSERVATION STATUS

Known from two collections within 1 – 2 km of each other at a single location, the higher altitudes of the Bali Ngemba Forest Reserve in NW Region.

*Lychnodiscus bali* is here assessed as Critically Endangered since only a single location is known, which is unprotected with clear threats of habitat clearance. The area of occupation is calculated as 4 km^2^ using the preferred IUCN grid cell size and the extent of occurrence is estimated as the same or slightly larger. Furthermore, less than 50 mature individuals have been observed. The first author saw only a single individual during several weeks surveying the forest (Cheek 10503). The species should be included in the next edition of the Red Data Book of Cameroon Plants (Onana & Cheek 2011).

Plotting the grid reference of the two specimens on Google Earth and examining historic imagery shows that there is an increase in cleared areas of forest likely due to the creation of more agricultural land by the local community which was observed by the first author when collecting the type specimen. This justifies an assessment of CR B1ab(iii) + B2ab(iii) + D. This distinctive and in flower conspicuous small tree has not been found in surveys elsewhere in the Cameroon Highlands and adjacent areas (Cheek 1992; Cheek *et al*. 1996; Cable & Cheek 1998; Cheek *et al*. 2000; Maisels *et al*. 2000; Chapman & Chapman 2001; Cheek *et al*. 2004; Harvey *et al*. 2004; Cheek *et al*. 2006; Cheek *et al*. 2010; Harvey *et al*. 2010; Cheek *et al*. 2011). Therefore, it may indeed be endemic to Bali Ngemba Forest Reserve, the only significant submontane forest surviving in the Bamenda Highlands which has seen > 95% loss of the original forest cover (Cheek *et al*. 2000; Murphy *et al*. 2022; 2023).

### ETYMOLOGY

Named as a noun in apposition for Bali Ngemba Forest Reserve, to which it appears globally unique.

### LOCAL NAMES & USES

None are known.

### NOTES

*Lychnodiscus bali* appears to be an ecological pioneer, benefitting from limited disturbance since both specimens were found in areas that had been previously cleared of forest for agriculture and were then regenerating as “farmbush” (*Cheek* 10503), or even with some crops still present (“beans and cocoyams” *Ghogue* 1080). The similar *Lychnodiscus grandifolius* also occurs in disturbed areas (Lachenaud et al. 2018). *Lychnodiscus bali* has the highest altitudinal range of any species in the genus (1700 – 1950 m alt.). We speculate that the species evolved from an ancestral species that occurred at lower altitudes as do all other species of the genus. The species may be pollinated by bees: “a lot of bees feeding on their white flowers (*Ghogue* 1080).

## Discussion

*Lychnodiscus bali* is the most recent of several new species of Sapindaceae to have been described from surveys in Cameroon, mainly from the Cameroon Highlands, other species having been *Allophylus ujori* Cheek (Cheek & Etuge 2009a), *Deinbollia oreophila* Cheek (Cheek & Etuge 2009b), *Allophylus bertoua* Cheek (Cheek & Haba 2016) and *Deinbollia onanae* Cheek (Cheek et al 2021). There is no doubt that additional new species remain to be discovered in Cameroon in this family which in tropical Africa is taxonomically under-researched.

The discovery from Bali Ngemba of *Lychnodiscus bali* is the latest of a steady stream of taxonomic novelties from this small, threatened remnant of submontane forest in the Bamenda Highlands. Other recently published species from Bali Ngemba have been *Vepris onanae* Cheek (Cheek et al. 2022b), *Monanthotaxis bali* Cheek (Cheek et al. 2022c) and *Deinbollia onanae* Cheek (Cheek *et al*. 2021). 34 threatened species are recorded from this forest, of which nine are strict endemics and 11 near-endemics (Cheek et al. 2022c).

The discovery of this new species among the specimens incorporated in the Kew Herbarium (K) supports previous evidence that the availability of, and ready access to, well-managed herbaria are important factors for completing our knowledge of the species on our planet (Bebber *et al*. 2010; Brown et al. 2023; Onana *et al*. 2017). However more trained taxonomists are needed to work on these collections, as well as in the field (Löbl et al. 2023; Sutcliffe & O’Reilly 2010).

*Lychnodiscus bali* is the latest in a series of discoveries of new species to science from the forest habitats of the Cameroon Highlands. These species vary from epiphytic herbs e.g. *Impatiens frithi* Cheek (Balsaminaceae, Cheek & Csiba 2002) and *I. etindensis* Cheek & Eb. Fisch. (Balsaminaceae, Cheek & Fischer 1999) to terrestrial herbs e.g. *Brachystephanus kupeensis* I. Darbysh. & Champl. (Acanthaceae, Champluvier & Darbyshire 2009) and *Isoglossa dispersa* I. Darbysh. (Darbyshire *et al*. 2011), rheophytes e.g. *Ledermanniella onanae* Cheek (Cheek 2003) and *Saxicolella ijim* Cheek (Podostemaceae; Cheek *et al*. 2022a) to achlorophyllous mycotrophs e.g. *Kupea martinetugei* Cheek & S.A.Williams (Triuridaceae, Cheek *et al*. 2003) and *Afrothismia amietii* Cheek (Cheek 2007, Afrothhismiaceae, Cheek et al. 2023) and understorey shrubs and small trees e.g. *Kupeantha kupensis* Cheek (Rubiaceae, Cheek *et al*. 2018a) and *Psychotria darwiniana* Cheek (Rubiaceae, Cheek *et al*. 2009), climbers e.g. *Sabicea bullata* Zemagho, O. Lachenaud & Sonké (Rubiaceae, Zemagho et al. 2014), *Ancistrocladus grandiflorus* Cheek (Ancistrocladaceae, Cheek 2000), to canopy trees, e.g. *Korupodendron songweanum* Litt & Cheek (Vochysiaceae, Litt & Cheek 2002), *Vepris montisbambutensis* Onana (Onana *et al*. 2015), *Vepris mbamensis* Onana (Onana *et al*. 2019), *Vepris zapfackii* Cheek & Onana (Cheek & Onana 2021) and *Carapa oreophila* Kenfack (Meliaceae, Kenfack 2011).

Most of these new species were first described as point or near-endemics, but several have, with more research, been found to be more widespread e.g. *Tricalysia elmar* Cheek (Rubiaceae, Cheek *et al*. 2020a) now known to extend from Mt Kupe and Bali-Ngemba to e.g. the Rumpi Hills (Lachenaud pers. obs. 2009), *Mussaenda epiphytica* Cheek (Rubiaceae, Cheek 2009), first only known from Bakossi, but then found from elsewhere (Lachenaud *et al*. 2013) and now found in Gabon (Lachenaud pers. obs.), *Coffea montekupensis* Stoff. (Stoffelen *et al*. 1997) first thought to be endemic to Mt Kupe is now known to occur in the Tofala Sanctuary (Lebialem Highlands, Harvey *et al*. 2010) and *Oxyanthus okuensis* Cheek & Sonké (Rubiaceae, Cheek & Sonké 2000) initially thought to be endemic to Mt Oku is now known to extend to Tchabal Mbabo (Lachenaud *et al*. 2013). It is to be hoped that the range of *Lychnodiscus bali* will be similarly extended by future research and its extinction risk assessment consequently reduced.

## Conclusion

It is important to uncover the existence of previously unknown plant species as soon as possible and to formally name them. Until this is done, they are invisible to science and the possibility of their being assessed for their conservation status and appearing on the IUCN Red List is greatly reduced (Cheek *et al*. 2020b), limiting the likelihood that they will be proposed for conservation measures, and that such measures will be accepted, affording protection. Completing our knowledge of the world’s plant species was target one of the Global Strategy for Plant Conservation, leading to assessments of their conservation status (target two) and in situ conservation as Important Plant Areas (target five), although meeting these has been challenging (Paton & Nic Lughadha 2010). Although there are exceptions (Cheek & Etuge 2009; Cheek *et al*. 2019a), most new plant species to science are highly range-restricted, making them almost automatically threatened (Cheek *et al*. 2020b).

Only 7.2% of the 369,000 flowering plant species (the number is disputed) known to science have been assessed on the IUCN Red List. (Bachman *et al*. 2019; Nic Lughadha *et al*. 2016; 2017). However, the vast majority of plant species still lack assessments on the Red List (Nic Lughadha *et al*. 2020). Fortunately, Cameroon has a plant Red Data book (Onana & Cheek 2011), which details 815 threatened species, but it needs updating. Thanks to the Global Tree Assessment (BGCI 2021) many of the world’s tree species have now been assessed. The State of the World’s Trees concluded that the highest proportion of threatened tree species is found in Tropical Africa, and that Cameroon has the highest number (414) of threatened tree species of all tropical African countries (BGCI 2021). This will be further increased by the addition of *Lychnodiscus bali*.

Concerns about global plant species extinctions are increasing as the biodiversity crisis continues. In Cameroon, the lowland forest species *Inversodicraea bosii* Cusset, *Oxygyne triandra* Schltr., *Afrothismia pachyantha* Schltr. have been considered extinct for some years (Cheek et al. 2017; Cheek & Williams 1999; Cheek *et al*. 2018b; Cheek *et al*. 2019b).

Similarly, *Pseudohydrosme bogneri* Cheek & Moxon-Holt and *P. buettneri* Engl. are now considered extinct in lowland forest in neighbouring Gabon (Moxon-Holt & Cheek 2020; Cheek *et al*. 2021b). However, Cameroonian endemic submontane (cloud) forest species are now also being recorded as globally extinct, such as the well-documented case of *Vepris bali* Cheek (Cheek *et al*. 2018c) also at Bali Ngemba Forest Reserve. Global species extinctions are being recorded from across Africa, from West (e.g. *Saxicolella deniseae* Cheek in Guinea (Cheek *et al*. 2022a) to East (e.g. *Cynometra longipedicellata* Harms, *Kihansia lovettii* Cheek and *Vepris* sp A of FTEA in Tanzania (Gereau et al. 2020; Cheek & Luke 2022; Cheek 2004).

If such extinctions are to cease, or more realistically, to be slowed, improved conservation prioritisation programmes are needed to firstly determine the most important plant areas for conservation (Darbyshire *et al*. 2017) and secondly to implement protection with local communities and authorities through well-drawn up management plans and resourcing. Cultivation and seedbanking of species at risk of extinction, if feasible, are important fall-back strategies, but conservation of species in their natural habitat must be the first priority.

The future of *Lychnodiscus bali* depends on greatly improved protection of the Bali Ngemba Forest Reserve which for years has seen steady encroachment, to the extent that at least one of its species is almost certainly globally extinct (*Vepris bali*) and another is feared likely to be extinct (*Monanthotaxis bali*, see above). It is hoped that recent listing of Bali Ngemba as an Important Plant Area (Murphy *et al*. 2022; 2023) will support better protection of this unique forest in future by local communities and administration.

## Supporting information

Supplemental file Holotype image

## Acknowledgements

This paper was completed as part of the Cameroon TIPAs (Tropical Important Plant Areas) project at RBG, Kew, which is supported by Players of Peoples Postcode Lottery. We thank Lydia Burns and Penny Appelbe of Kew’s Foundation for making this possible. The second author’s contribution to this paper was made possible by visits from Cameroon to RBG, Kew, U.K. sponsored by the Bentham-Moxon Trust of RBG, Kew. The specimens cited in this paper were collected with the support of volunteers of Earthwatch Europe, Oxford and by our colleagues including Kenneth Tah, Olivier Sene, Victor Nana, Verina Ingram, David Okebiro, Assefa, B. Gapta, H. Ndue, M. Kissimou, Rene Nfon, Stuart Cable, Ben Pollard and the late Martin Etuge. This paper is a result of the partnership between RBG, Kew and IRAD-National Herbarium of Cameroon, Yaoundé and we thank the late Dr Benoît Satabié, Drs Gaston Achoundong, Florence Ngo Ngwe, Eric Nana, Jean Lagarde Betti, the current and former directors or acting Directors, of IRAD-National Herbarium of Cameroon, Yaoundé, for expediting the collaboration between our two institutes. Dr Jean Paul Ghogue is thanked for collecting one of the specimens cited. Two anonymous reviewers are thanked for constructively reviewing an earlier version of this paper.

This paper is dedicated to the late Janis Carole Shillito (1941-2023) who typed this and many other manuscripts on the taxonomy and conservation of plants, especially those of Cameroon, assisting the first author, over more than 10 years.

The authors declare that they have no conflict of interest.

## References

African Plant Database (version 3.4.0). (continuously updated). Conservatoire et Jardin botaniques de la Ville de Genève and South African National Biodiversity Institute, Pretoria, Retrieved August 2022 from http://www.ville-ge.ch/musinfo/bd/cjb/africa/

Bachman, S.P., Field, R., Reader, T., Raimondo, D., Donaldson, J., Schatz, G.E. and Lughadha, E.N. (2019). Progress, challenges and opportunities for Red Listing. Biological Conservation 234: 45–55. 10.1016/j.biocon.2019.03.002

Bates, L. (2005). Cognitive aspects of travel and food location by chimpanzees (Pan troglodytes schweinfurthii) of the Budongo forest reserve, Uganda. PhD thesis, University of St Andrews.

Bebber, D.P., Carine, M.A., Wood, J.R.I., Wortley, A.H., Harris, D.J., Prance, G.T., Davidse, G., Paige, J., Pennington, T.G., Robson, N.K.B. & Scotland, R.W. (2010). Herbaria are a major frontier for species discovery. Proceedings of the National Academy of Sciences of the U.S.A. http://www.pnas.org/cgi/contens/short/1011841108

Beentje, H. & Cheek, M. (2003). Glossary. In Beentje, (ed.), Flora of Tropical East Africa. Balkema, Lisse, Netherlands.

BGCI (2021). State of the World’s Trees. Botanic Gardens Conservation International (BGCI), Richmond, UK.

Bosch, C. H. (2012). Lychnodiscus cerospermus p. 444 – 445 in Lemmens, R. H. M. J., Louppe, D., & Oteng-Amoako, A. A. Timbers 2 (Vol. 7). PROTA.

Brown M.J.M., Bachman S.P. and Nic Lughadha E. (2023). Three in four undescribed plant species are threatened with extinction. Letter. New Phytologist http://www.newphytologist.com

Buerki, S., Callmander, M.W., Acevedo-Rodriguez, P., Lowry, P.P., Munzinger, J., Bailey, P., Maurin, O., Brewer, G.E., Epitawalage, N., Baker, W.J. and Forest, F., 2021. An updated infra-familial classification of Sapindaceae based on targeted enrichment data. American Journal of Botany, 108(7), pp.1234 – 1251.

Burkill, H.M. (2000). The Useful Plants of West Tropical Africa. Vol. 5, families S-Z. Royal Botanic Gardens, Kew.

Cable, S. & Cheek, M. (1998). The Plants of Mt Cameroon, a Conservation Checklist. Royal Botanic Gardens, Kew.

Champluvier, D. & Darbyshire, I. (2009). A revision of the genera Brachystephanus and Oreacanthus (Acanthaceae) in tropical Africa. Systematics and Geography of Plants 79(2):115–192. 10.2307/25746

Chapman, J. & Chapman, H. (2001). The Forests of Taraba and Adamawa States, Nigeria an Ecological Account and Plant Species Checklist. University of Canterbury: Christchurch, New Zealand. pp. 221.

Cheek, M. (1992). A Botanical Inventory of the Mabeta-Moliwe Forest. Royal Botanic Gardens, Kew.

Cheek, M. (2003). A new species of Ledermanniella (Podostemaceae) from western Cameroon. Kew Bulletin 58: 733–737. 10.2307/4111153

Cheek, M. (2007). Afrothismia amietii (Burmanniaceae), a new species from Cameroon. Kew Bull. 61: 605–607. http://www.jstor.org/stable/20443306

Cheek, M. (2009). Mussaenda epiphytica sp. nov. (Rubiaceae_) an epiphytic shrub from cloud forest of the Bakossi Mts, western Cameroon. Nordic J. Bot. 27(6): 456–459. 10.1111/j.1756-1051.2009.00576.x

Cheek, M. (2017). Microcos magnifica (Sparrmanniaceae) a new species of cloudforest tree from Cameroon. PeerJ 5:e4137 10.7717/peerj.4137

Cheek, M. (2023). Identification and Naming pp 100–103 in Davies, N.M.J., Drinkell, C, Utteridge, T.M.A. (Eds.) The Herbarium Handbook. Kew Publishing

Cheek, M. & Cable, S. (1997). Plant Inventory for conservation management: the Kew-Earthwatch programme in Western Cameroon, 1993 – 96, pp. 29–38 in Doolan, S. (Ed.) African Rainforests and the Conservation of Biodiversity, Earthwatch Europe, Oxford.

Cheek, M. & Csiba, L. (2002). A new epiphytic species of Impatiens (Balsaminaceae) from western Cameroon. Kew Bull. 57: 669 – 674. 10.2307/4110997

Cheek, M. & Etuge, M. (2009). Allophylus conraui (Sapindaceae) reassessed and Allophylus ujori described from the Cameroon Highlands of West Africa. Kew Bull. 64: 495–502. 10.1007/s12225-009-9139-x

Cheek, M. & Etuge, M. (2009). A new submontane species of Deinbollia (Sapindaceae) from Western Cameroon and adjoining Nigeria. Kew Bull. 64: 503–508. 10.1007/s12225-009-9132-4

Cheek, M. & Fischer, E. (1999). A tuberous and epiphytic new species of Impatiens (Balsaminaceae) from Southwest Cameroon. Kew Bull. 54: 471–475. 10.2307/4115828

Cheek, M. & Haba, P. M. (2016). Spiny African Allophylus (Sapindaceae): a synopsis. Kew Bulletin 71: 55. 10.1007/S12225-016-9672-3

Cheek, M. & Luke, W.R.Q. (2022). A taxonomic synopsis of unifoliolate continental African Vepris (Rutaceae) with three new threatened forest tree species from Kenya and Tanzania and two possibly extinct. Kew Bull. 1–29. 10.1007/s12225-023-10120-0

Cheek, M. & Onana, J.M. (2021). The endemic plant species of Mt Kupe, Cameroon with a new Critically Endangered cloud-forest tree species, Vepris zapfackii (Rutaceae). Kew Bull 76, 721–734 10.1007/s12225-021-09984-x

Cheek, M. & Sonké, B. (2000). A new species of Oxyanthus (Rubiaceae-Gardeniinae) from western Cameroon. Kew Bulletin 55: 889–893. 10.2307/4113634

Cheek, M. & Williams, S. (1999). A Review of African Saprophytic Flowering Plants. In: Timberlake, Kativu eds. African Plants. Biodiversity, Taxonomy & Uses. Proceedings of the 15th AETFAT Congress at Harare. Zimbabwe, 39–49.

Cheek, M., Achoundong, G., Onana, J-M., Pollard, B., Gosline, G., Moat, J., Harvey, Y.B. (2006). Conservation of the Plant Diversity of Western Cameroon. In: Ghazanfar SA, H.J. Beentje (eds). Proceedings of the 17th AETFAT Congress, Addis Ababa. kEthiopia, 779–791.

Cheek, M., Alvarez-Agiurre, M.G., Grall, A., Sonké, B., Howes, M-J.R., Larridon, I. (2018a). Kupeantha (Coffeeae, Rubiaceae), a new genus from Cameroon and Equatorial Guinea. PLoS ONE 13: 20199324. 10.1371/journal.pone.0199324

Cheek, M., Cable, S., Hepper, F.N., Ndam, N., Watts, J. (1996). Mapping plant biodiversity on Mt. Cameroon. In: Maesen, Burgt Rooy eds. The Biodiversity of African Plants (Proceedings XIV AETFAT Congress. Cameroon: Kluwer, 110–120. 10.1007/978-94-009-0285-5_16

Cheek, M., Causon, I., Tchiengue, B. & House, E. (2020a). Notes on the endemic cloud forest plants of the Cameroon Highlands and the new, Endangered, Tricalysia elmar (Coffeeae-Rubiaceae). Plant Ecology and Evolution 153: 167–176 10.5091/plecevo.2020.1661

Cheek, M., Corcoran, M. & Horwath, A. (2009b). Four new submontane species of Psychotria (Rubiaceae) with bacterial nodules from western Cameroon. Kew Bull. 63: 405 – 418. 10.1007/s12225-008-9056-4

Cheek, M., Darbyshire, I. & Onana, J.-M. (2022c). Monanthotaxis bali (Annonaceae) a new Critically Endangered (possibly extinct) montane forest treelet from Bali Ngemba, Cameroon. BioRxiv 10.1101/2022.07.04.498636

Cheek, M., Etuge, M. & Williams, S. (2019b). Afrothismia kupensis sp. nov. (Thismiaceae), Critically Endangered, with observations on its pollination and notes on the endemics of Mt Kupe, Cameroon. Blumea 64: 158 – 164. 10.3767/blumea.2019.64.02.06

Cheek M, Feika A, Lebbie A, Goyder D, Tchiengue B, Sene O, P. Tchouto P, Burgt X. (2017). A synoptic revision of Inversodicraea (Podostemaceae). Blumea 62, 2017: 125–156. 10.3767/blumea.2017.62.02.07

Cheek, M., Gosline, G. & Onana, J.M. (2018c). Vepris bali (Rutaceae), a new critically endangered (possibly extinct) cloud forest tree species from Bali Ngemba, Cameroon. Willdenowia 48: 285–292. 10.3372/wi.48.48207

Cheek, M., Harvey, Y.B., Onana, J-M. (2010). The Plants of Dom. Bamenda Highlands, Cameroon: A Conservation Checklist. Royal Botanic Gardens, Kew.

Cheek M, Harvey Y, Onana J-M. (2011). The Plants of Mefou Proposed National Park. Yaoundé, Cameroon: A Conservation Checklist. Royal Botanic Gardens, Kew.

Cheek, M., Hatt, S., & Onana, J. M. (2022b). Vepris onanae (Rutaceae), a new Critically Endangered cloud-forest tree species, and the endemic plant species of Bali Ngemba Forest Reserve, Bamenda Highlands Cameroon. Kew Bull., 1–15. 10.1007/s12225-022-10020-9

Cheek, M., Molmou, D., Magassouba, S., & Ghogue, J. P. (2022a). Taxonomic revision of Saxicolella (Podostemaceae), African waterfall plants highly threatened by Hydro-Electric projects. Kew Bull., 1 – 31. 10.1007/s12225-022-10019-2

Cheek, M., Nic Lughadha, E., Kirk, P., Lindon, H., Carretero, J., Looney, B., Douglas, B., Haelewaters, D., Gaya, E., Llewellyn, T., Ainsworth, M., Gafforov, Y., Hyde, K., Crous, P., Hughes, M., Walker, B.E., Forzza, R.C., Wong, K.M., Niskanen, T. (2020b). New scientific discoveries: plants and fungi. Plants, People Planet 2: 371 – 388. 10.1002/ppp3.10148 https://doi.org/10.7717/peerj.11036

Cheek, M., Onana, J.M., Chapman, H.M. (2021a). The montane trees of the Cameroon Highlands, West-Central Africa, with Deinbollia onanae sp. nov. (Sapindaceae), a new primate-dispersed, Endangered species. PeerJ 9:e11036

Cheek, M., Onana, J-M., Pollard, B.J. (2000). The Plants of Mount Oku and the Ijim Ridge, Cameroon, a Conservation Checklist. Royal Botanic Gardens, Kew.

Cheek, M., Onana, J-M., Yasuda, S., Lawrence, P., Ameka, G., Buinovskaja G. (2019a). Addressing the Vepris verdoorniana complex (Rutaceae) in West Africa, with two new species. Kew Bull. 74: 53. 10.1007/S12225-019-9837-Y

Cheek, M., Pollard, B.J., Darbyshire, I, Onana, J.M. & Wild, C. (2004). The Plants of Kupe, Mwanenguba and the Bakossi Mts, Cameroon. A Conservation Checklist. Royal Botanic Gardens, Kew.

Cheek, M., Soto Gomez, M., Graham, S.W. & Rudall, P.J. (2023). Afrothismiaceae (Dioscoreales), a new fully mycoheterotrophic family endemic to tropical Africa. bioRxiv 10.1101/2023.01.10.523343

Cheek, M., Tchiengue, B., Tacham, W.N. (2017). Ternstroemia cameroonensis (Ternstroemiaceae), a new medicinally important species of montane tree, nearly extinct in the Highlands of Cameroon. Blumea 62: 53 – 57.10.3767/000651917X695362.

Cheek, M., Tchiengué, B., van der Burgt, X. (2021b). Taxonomic revision of the threatened African genus Pseudohydrosme Engl. (Araceae), with P. ebo, a new, critically endangered species from Ebo, Cameroon. PeerJ 9:e10689 10.7717/peerj.10689.

Cheek, M., Tsukaya, H., Rudall, P.J., Suetsugu, K. (2018b). Taxonomic monograph of Oxygyne (Thismiaceae), rare achlorophyllous mycoheterotrophs with strongly disjunct distribution. PeerJ 6: e4828. 10.7717/peerj.4828

Cheek, M., Williams, S. & Etuge, M. (2003). Kupea martinetugei, a new genus and species of Triuridaceae from western Cameroon. Kew Bulletin 58: 225–228. 10.2307/4119366

Darbyshire, I., Anderson, S., Asatryan, A., Byfield, A., Cheek, M., Clubbe, C., Ghrabi, Z., Harris, T., Heatubun, C. D., Kalema, J., Magassouba, S., McCarthy, B., Milliken, W., Montmollin, B. de, Nic Lughadha, E., Onana, J.M., Saidou, D., Sarbu, A., Shrestha, K. & Radford, E. A. (2017). Important Plant Areas: revised selection criteria for a global approach to plant conservation. Biodivers. Conserv. 26: 1767–1800. 10.1007/s10531-017-1336-6.

Darbyshire, I., Pearce, L. & Banks, H. (2011). The genus Isoglossa (Acanthaceae) in west Africa. Kew Bulletin 66 (3): 425–439. 10.1007/s12225-011-9292-x

Davies, F.G. & Verdcourt, B. (1998). Sapindaceae. Flora of Tropical East Africa. Rotterdam: Balkema.

Eilu, G., Hafashimana, D. L., & Kasenene, J. M. (2004). Tree species distribution in forests of the Albertine Rift, western Uganda. African Journal of Ecology, 42(2), 100 – 110.

Fouilloy, R. & Hallé, N. (1973). Sapindacées. Flore du Cameroun 16. Paris: Museum National d’Histoire Naturelle.

Gereau, R.E., Kabuye, C., Kalema, J., Kamau, P., Kindeketa, W., Luke, W.R.Q., Lyaruu, H.V.M., Malombe, I., Mboya, E.I., Mollel, N., Njau, E.-F., Schatz, G.E., Sitoni, D., Ssegawa, P. & Wabuyele, E. (2020). Cynometra longipedicellata. The IUCN Red List of Threatened Species 2020: e.T32275A2812477. 10.2305/IUCN.UK.2020-3.RLTS.T32275A2812477.en. Downloaded on 01 May 2022.

Gosline, G., Bidault, E., van der Burgt, X. et al. (2023a). A Taxonomically-verified and Vouchered Checklist of the Vascular Plants of the Republic of Guinea. Sci Data 10, 327 (2023). 10.1038/s41597-023-02236-6

Gosline, G. et al. (2023b). Checklist of the Vascular Plants of the Republic of Guinea– printable format (1.10). Zenodo. 10.5281/zenodo.7734985

Hauman, L. (1960). Sapindaceae. In: Robyns, W., Staner, P., Demaret, F., Germain, R., Gilbert, G., Hauman, L., Homès, M., Jurion, F., Lebrun, J., Vanden Abeele, M. & Boutique, R. (Editors). Flore du Congo Belge et du Ruanda-Urundi. Spermatophytes. Volume 9. Institut National pour l’Étude Agronomique du Congo belge, Brussels, Belgium. pp. 279– 384.

Harvey, Y.B., Pollard, B.J., Darbyshire, I., Onana, J.-M., Cheek, M. (2004). The Plants of Bali Ngemba Forest Reserve. Cameroon: A Conservation Checklist. Royal Botanic Gardens, Kew.

Harvey, Y.B., Tchiengue, B., Cheek, M. (2010). The Plants of the Lebialem Highlands, a Conservation Checklist. Royal Botanic Gardens, Kew.

IPNI (continuously updated). The International Plant Names Index. http://ipni.org/.

IUCN. 2012. IUCN Red List Categories and Criteria: Version 3.1. Second edition. Gland, Switzerland and Cambridge, UK: IUCN. Available from: http://www.iucnredlist.org/ x(accessed: 01/2017).

Joyce, E.M., Appelhans, M.S., Buerki, S., Cheek, M., de Vos, J.M., Pirani, J.R., Zuntini, A.R., Bachelier, J.B., Bayly, M.J., Callmander, M.W. and Devecchi, M.F. (2023). Phylogenomic analyses of Sapindales support new family relationships, rapid Mid-Cretaceous Hothouse diversification, and heterogeneous histories of gene duplication. Frontiers in Plant Science 14:1063174. 10.3389/fpls.2023.1063174

JStor Global Plants. (continuously updated), Available at http://plants.jstor.org/(accessed 14 June 2022).

Keay, R.W.J. (1958). Sapindaceae, pp.709–725 in Keay (ed.) Flora of West Tropical Africa 1(2). London: Crown Agents.

Kenfack D. (2011). A synoptic revision of Carapa (Meliaceae). Harvard Papers in Botany 16(2), 171–231. http://www.jstor.org/stable/41761712

Lachenaud, O., Droissart, V., Dessein, S., Stévart, T., Simo, M., Lemaire, B., Taedoumg, H. and Sonké, B. (2013). New records for the flora of Cameroon, including a new species of Psychotria (Rubiaceae) and range extensions for some rare species. Plant Ecology and Evolution 146: 121 – 133. 10.5091/plecevo.2013.632

Lachenaud, O., Stévart, T., Boupoya, A., Texier, N., Dauby, G., & Bidault, E. (2018). Novitates Gabonenses 88: additions to the flora of Gabon and new records of little-known species. Plant Ecology and Evolution, 151(3), 393–422.

Lebrun, J.-P. and Stork, A.L. (1992). Enumération des plantes à fleurs d’Afrique tropicale. Vol. II : Chrysobalanaceae à Apiaceae. Genève: Conservatoire et Jardin Botaniques.

Löbl, I., Klausnitzer, B.,Hartmann, M., Krell, F.-T. (2023) The Silent Extinction of Species and Taxonomists—An Appeal to Science Policymakers and Legislators. Diversity, 15: 1053. 10.3390/d15101053

Maisels, F.M., Cheek, M., Wild, C. (2000). Rare plants on Mt Oku summit, Cameroon. Oryx 34: 136 – 140. 10.1017/s0030605300031057.

Moxon-Holt, L. & Cheek, M. (2020). Pseudohydrosme bogneri sp. nov. (Araceae), a spectacular Critically Endangered (Possibly Extinct) species from Gabon, long confused with Anchomanes nigritianus. BioRxiv (pre-print) 10.1101/2021.03.25.437040

Murphy, B, Tah, K, Cheek, M., Njoya, M. (2022). Tropical Important Plant Areas Explorer: Bali Ngemba Forest (Cameroon). https://tipas.kew.org/site/bali-ngemba-forest-reserve-2/ (Accessed on 08/10/2022)

Murphy, B., Onana, J.M.. van der Burgt, X. M., Tchatchouang Ngansop, E., Williams, J., Tchiengué, B., Cheek, M. (2023). Important Plant Areas of Cameroon. Royal Botanic Gardens, Kew.

Nic Lughadha, E., Bachman, S.P., Govaerts, R. (2017). Plant fates and states: Response to Pimm & Raven. Trends in Ecology and Evolution 32: 887–889

Nic Lughadha, E., Bachman, S.P., Leão, T.C., Forest, F., Halley, J.M., Moat, J., Acedo, C., Bacon, K.L., Brewer, R.F., Gâteblé, G., Gonçalves, S.C., Govaerts, R., Hollingsworth, P.M., Krisai-Greilhuber, I., de Lirio, E.J., Moore, P.G.P., Negrão, R., Onana, J.M., Rajaovelona, L.R., Razanajatovo, H., Reich, P.B., Richards, S.L., Rivers, M.C., Amanda Cooper, A., Iganci, J., Lewis, G.L., Smidt, E.C., Antonelli, A., Mueller, G.M. & Walker, B.E. (2020). Extinction risk and threats to plants and fungi. Plants, People, Planet 2: 389–408. 10.1098/rstb.2017.0402

Nic Lughadha, E., Govaerts, R., Belyaeva, I., Black, N., Lindon, H. Allkin, R. Magill R.E. & Nicolson. (2016). Counting counts: Revised estimates of numbers of accepted species of flowering plants, seed plants, vascular plants and land plants with a review of other recent estimates. Phytotaxa 272: 82–88. 10.11646/phytotaxa.272.1.5

Onana, J.-M. & Cheek, M. (2011). The Red Data Book, Plants, of Cameroon.Royal Botanic Gardens, Kew

Onana J. M. & Chevillotte H. (2015). Taxonomie des Rutaceae-Toddalieae du Cameroun revisitée : découverte de quatre espèces nouvelles, validation d’une combinaison nouvelle et véritable identité de deux autres espèces de Vepris Comm. ex A.Juss. Adansonia, sér. 3, 37 (1): 103–129. 10.5252/a2015n1a7

Onana J. M., Cheek M. & Chevillotte H. (2019). Additions au genre Vepris Comm. ex A.Juss. (Rutaceae-Toddalieae) au Cameroun. Adansonia, sér. 3, 41 (5): 41–52. 10.5252/adansonia2019v41a5 http://adansonia.com/41/5

Onana J.M., Mbome M. J. & Mekembom, Y.N (2017) The north-south synergy: the National Herbarium and Limbe Botanic Garden experience In: Friis, I.and Balslev, H. (eds) Proceedings of an international symposium held by The Royal Danish Academy of Sciences and Letters in Copenhagen, 19th–21st of May, 2015. Scientia Danica. Series B, Biologica vol. 6: 117– 139.

Paton, A., & Nic Lughadha, E.M. (2011). The irresistible target meets the unachievable objective: what have 8 years of GSPC implementation taught us about target setting and achievable objectives? Botanical Journal of the Linnean Society 166(3): 250–260.

POWO (continuously updated). Plants of the World Online. Facilitated by the Royal Botanic Gardens, Kew. http://www.plantsoftheworldonline.org/ xRetrieved 28 Aug. 2022.

Radlkofer, L. (1932). Sapindaceae in Engler, A. Das Pflanzenreich IV. 165 Heft 98c. Berlin: Wilhelm Engelmann.

Sosef, M.S.M., Wieringa, J.J., Jongkind, C.C.H., Achoundong, G., Azizet Issembé, Y., Bedigian, D., Van Den Berg, R.G., Breteler, F.J., Cheek, M., Degreef, J. (2006). Checklist of Gabonese Vascular Plants.Scripta Botanica Belgica 35. National Botanic Garden of Belgium. 435 pp.

Stoffelen, P., Cheek, M., Bridson, D., Robbrecht, E. (1997.) A new species of Coffea (Rubiaceae) and notes on Mt Kupe (Cameroon). Kew Bulletin 52(4): 989–994. 10.2307/3668527

Sutcliffe, J. & O’Reilly, C. (2010). Ecological skills: Mind the gap(s). Kew Bulletin 65: 529– 538

Thiers, B. (continuously updated). Index Herbariorum: A global directory of public herbaria and associated staff. New York Botanical Garden’s Virtual Herbarium. [continuously updated]. Available from: http://sweetgum.nybg.org/ih/(accessed: July 2022).

Zemagho, L.A., Lachenaud, O., Dessein, S. Liede-Schumann, S. & Sonké, B. (2014). Two new Sabicea (Rubiaceae) species from West Central Africa: Sabicea bullata and Sabicea urniformis. Phytotaxa 173 (4): 285–292. https://www.biotaxa.org/Phytotaxa/article/view/phytotaxa.173.4.3

