## Supplementary figures and images for "A new cloud forest tree *Lychnodiscus bali* (Sapindaceae), Critically Endangered from the Bali-Ngemba Forest reserve, N.W. Region, Cameroon"

### Supplemental file Holotype image

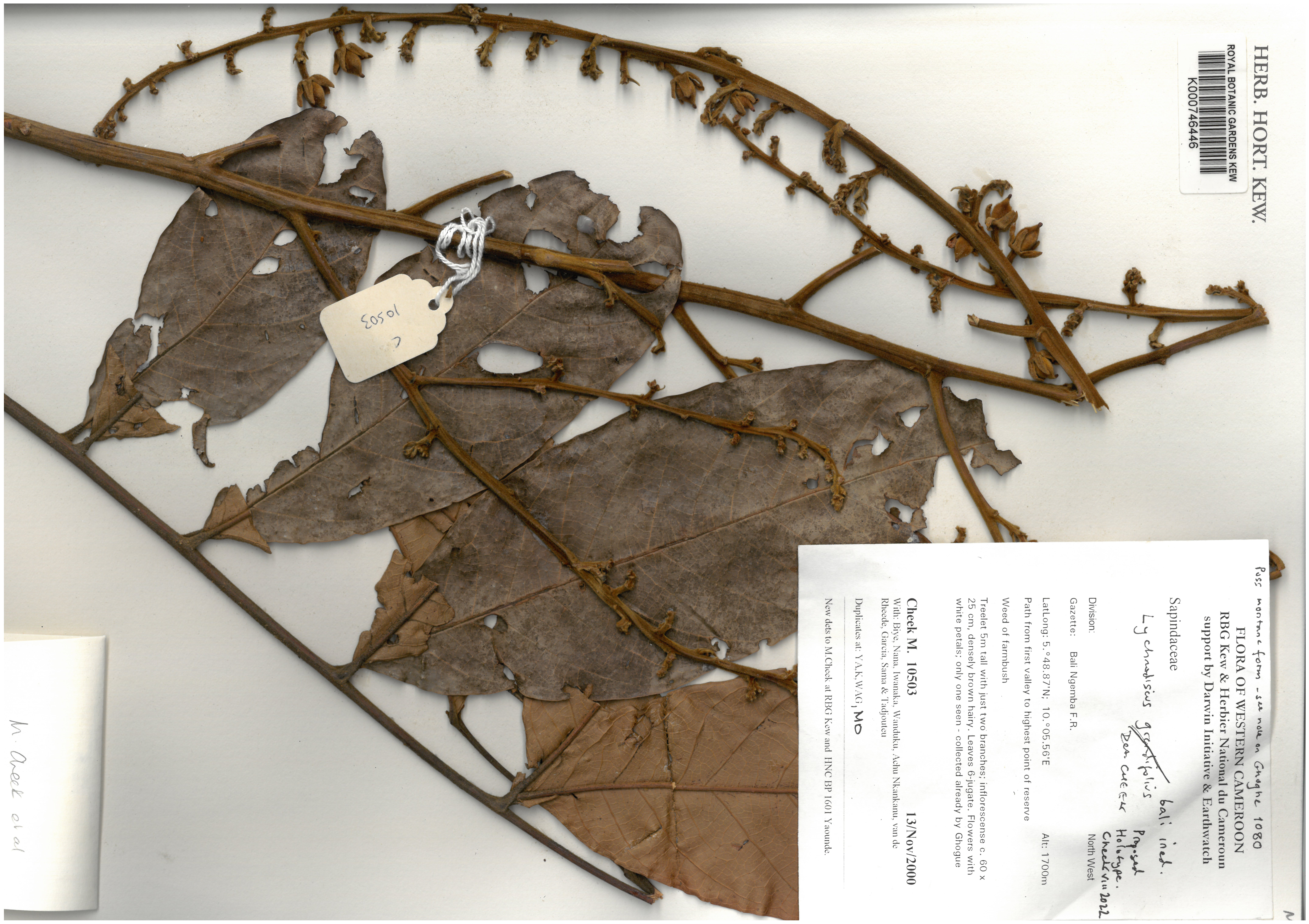
